# Goal-directed hippocampal theta sweeps during memory-guided navigation

**DOI:** 10.1101/2025.08.26.672489

**Authors:** Wenbo Tang, Xiyu Mei, Ryan E. Harvey, Estrella Carbajal-Leon, Talia Netzer, Hongyu Chang, Azahara Oliva, Antonio Fernandez-Ruiz

**Affiliations:** Department of Neurobiology and Behavior, Cornell University, Ithaca, NY, USA

## Abstract

During navigation, animals continually sample their surrounding space and plan routes to distant goals. The brain mechanisms underlying these behaviors and how they coordinate to support memory-guided navigation in open environments are not understood. Using large-scale recordings in rats, we found two distinct types of place cell sequences within theta cycles that encoded trajectories sweeping beyond the animal’s location: stereotypic left-right alternating sweeps and learning-dependent goal-directed sweeps. Goal-directed sweeps predicted upcoming trajectories to remembered goal locations, were coordinated with prefrontal cortex activity and preferentially replayed during sharp-wave ripples. We further describe a circuit mechanism in which a subpopulation of CA1 cells encodes egocentric goal direction, combined with reduced feedback inhibition, to generate goal-directed theta sweeps. These results indicate a flexible mechanism to support different behavioral demands during navigation.

## Main text

Animals explore their environment while constantly sampling their surroundings. They also navigate in a goal-directed manner to remembered locations such as food sources or shelters. Different neural mechanisms have been proposed to support these behaviors, with the hippocampal formation playing a central role (*1–4*). Hippocampal *place cells* fire at specific locations and generate a cognitive map of the environment as a population (*1*). In addition, place cells’ firing is coordinated by the hippocampal theta rhythm, generating compressed temporal sequences encoding spatial trajectories that start at locations behind the animal and extend ahead of it (“theta sequences” or “sweeps” (*5*–*7*)). These theta sequences have been profusely studied in rodents navigating linear mazes and found to sweep ahead of the path followed by the animal and to alternate between possible maze arms or goal locations at choice points, leading to the proposal that they represent possible behavioral alternatives to support goal-directed navigation (*8–13*). An alternative possibility is that theta sequences do not reflect cognitive processes such as trajectory planning or deliberation, but instead arise rigidly from intrinsic circuit dynamics and locomotion variables (*6, 14–16*). Distinguishing between these alternatives and determining the role of theta sequences in trajectory planning would require examining them during more naturalistic 2D navigation tasks, where animal movements are not constrained to linear tracks and decisions are not limited to binary arm choices. A recent study analyzed theta sequences in rats free foraging in open arenas and reported that, instead of sweeping forward along the travelled path, they alternated stereotypically in left and right lateral sweeps ahead of the animal (*14*). In contrast to previous claims, this evidence suggests that theta sweeps are an experience-independent mechanism to sample the surrounding environment, regardless of goals and cognitive demands (*14*). However, the dynamics of theta sweeps during unconstrained memory-guided navigation and how they are modulated by learning remain to be examined. To determine whether theta sequences are a mechanism for planning goal-directed trajectories, unbiased spatial sampling, or both, it is necessary to analyze them during both, free foraging as well as goal-directed navigation in open arenas.

If theta sequences support trajectory planning in open environments, they would need to be coupled with a vectorial signal that indicates the direction of the goal relative to the animal’s position (in egocentric coordinates), even when the goal is at a distant location. Such sequences should anticipate upcoming chosen trajectories regardless of the animal’s current heading. Furthermore, since goal-directed navigation relies not only on the hippocampus but also on its interactions with multiple brain areas, including the prefrontal cortex (PFC; (*17–21*)), these brain areas involved in goal-directed navigation and planning would be expected to coordinate with the hippocampus during goal-directed theta sweeps. To tackle these questions, we simultaneously recorded hundreds of neurons in the hippocampus and PFC of rats during random foraging and a goal-directed navigation learning task in an open maze.

## Results

### Lateral alternating and goal-directed hippocampal theta sweeps during foraging

We performed large-scale neural ensemble recordings using ultra-high density silicon probes in the dorsal hippocampal CA1 region and the PFC (pre-limbic cortex) of rats. Animals (*n* = 7) were trained to perform two different tasks in the same “cheeseboard maze” (*13, 22–24*): random foraging (*rf*) and goal-directed navigation (*gdn*) to three hidden rewarded locations that changed in every session (Fig. 1A). Simultaneous recordings of hundreds of neurons allowed us to analyze hippocampal theta sequences in 2D during individual oscillation cycles (fig. S1). We used a Bayesian decoding approach to analyze spatial representations in theta sequences. During random foraging, rats explored the maze to search for randomly baited water rewards. During animal navigation, we observed that theta sequences swept left and right ahead of the rat, alternating direction in successive theta cycles (Fig. 1B). These theta sweeps had an average angle of ∽30° with respect to the animal heading direction (Fig. 1, B-D) and an average length of 22.0 ± 2.5 cm (mean ± SD; fig. S2). Lateral alternating sweeps dominated theta sequence dynamics in all animals during random foraging sessions (Fig. 1, D and E and fig. S2), in agreement with previous reports (*14, 15*).

**Fig. 1.**
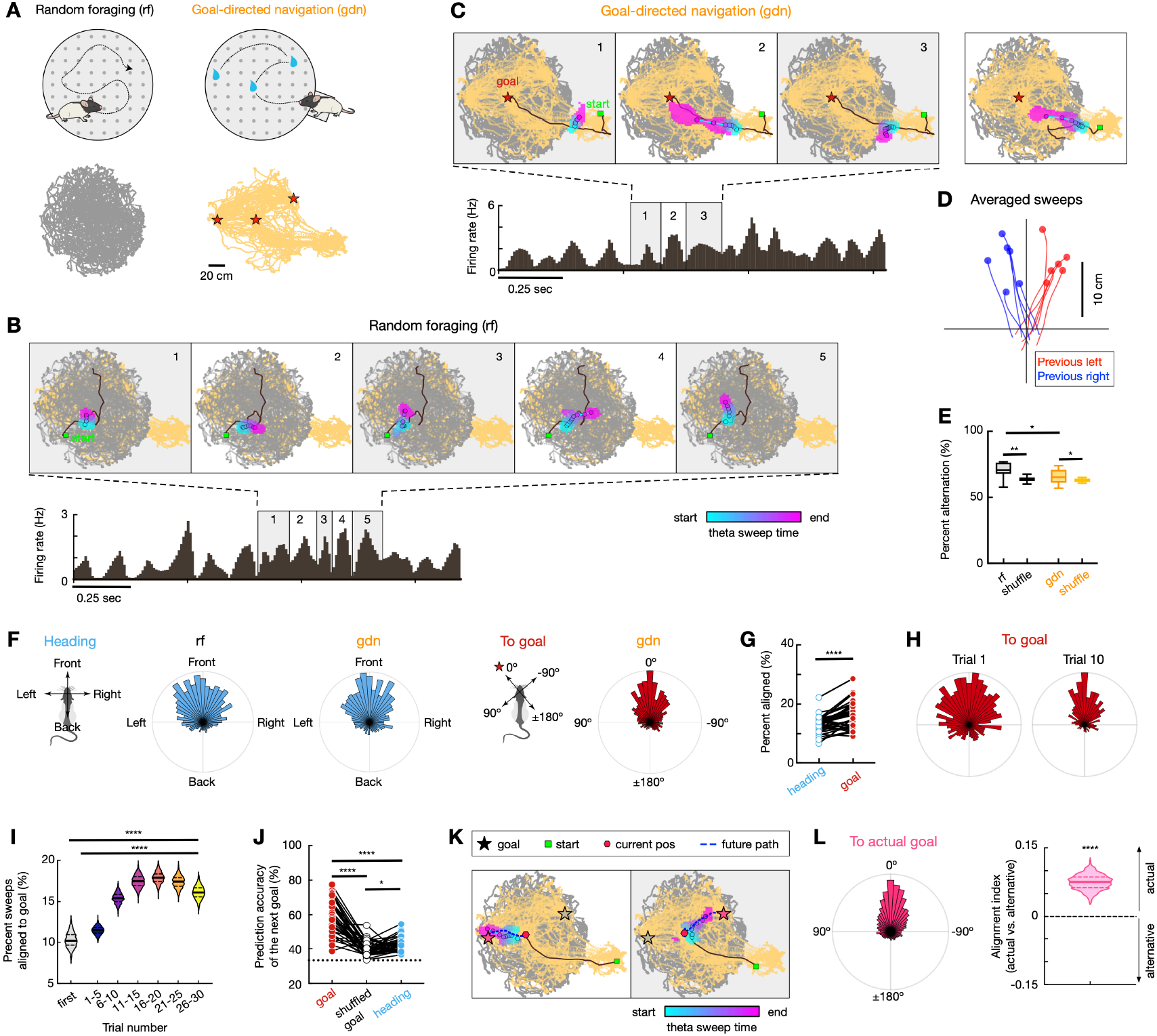
Theta sweeps alternate laterally during random foraging and become goal-directed during memory-guided navigation. **(A)** Cheeseboard maze diagram, showing random foraging (*rf*) and goal-directed navigation (*gdn*) tasks. Example animal trajectories are shown below. **(B)** Left–right alternating theta sweeps in CA1 during *rf. Top*: Sweeps decoded from CA1 population activity over five successive theta cycles (gray and yellow traces: positions visited during *rf* and *gdn*, respectively; black lines: current trajectory; green square: trajectory start; colored blobs: decoded positions, colored by time within sweep). *Bottom*: CA1 averaged firing rate showing 8–10 Hz theta modulation. **(C)** Goal-directed theta sweeps during *gdn* (data presented as in **B**). First and third sweeps were lateral; the second sweep points to the upcoming goal (star). An additional goal-directed sweep example that does not align with the heading is shown on the right. **(D)** Sweeps rotated to head-centered coordinates and averaged across theta cycles in which the preceding sweep went left (red) or right (blue). Data from 6 example sessions. **(E)** Triplet sweep patterns (L–R–L or R–L–R) occurred more frequently than shuffled controls (*rf*: *n* = 12 sessions, 6 rats; *gdn*: *n* = 63 sessions, 5 rats; \*\**p* = 0.0078, \**p* = 0.032, Wilcoxon paired test; \**p*(*rf* vs. *gdn*) = 0.029, rank-sum test). **(F)** Distributions of theta sweep directions relative to heading (*left*) and next goal chosen (*right*) from the example session shown in **A-C. (G)** A greater percentage of theta sweeps aligned (≤ 10º) to the goal direction than to the heading direction (\*\*\**p* = 1.33e-5, Wilcoxon paired test). **(H)** Goal-centered sweep direction distributions during trial 1 vs. trial 10 across all sessions. **(I)** Fraction of goal-directed sweeps across trials (\*\*\**p* < 1e-16, one-way ANOVA with *post hoc* test; bootstrapped *n* = 100). **(J)** Theta sweep directions predicted better the next goal chosen than heading direction or shuffled goals (\*\*\*\**p* = 3.29e-11, \**p* = 0.028, one-way repeated-measures ANOVA). **(K)** Examples of theta sweeps predicting the upcoming goal (red hexagon: current position; blue dashed line: future trajectory; pink star: chosen goal; gray star: alternative goal). **(L)** When the heading was biased toward a not chosen goal, theta sweep alignment to the chosen goal was still stronger than to the alternative goal (\*\*\*\**p* = 6.01e-6, bootstrap test). *N* = 36 sessions (from 5 rats; see Methods) in panels **F–L**.

In the goal-directed navigation task, rats had to discover three new hidden water reward locations in each session. They run 20-30 successive trials starting from a home box adjacent to the maze, find the three rewards, and returning to the box (*13, 24*). All animals showed consistent learning performance as evidenced by the decreased latency and path length to retrieve all rewards across trials (fig. S1C). In this task, we observed lateral theta sweeps alternating between left and right directions during navigation, although to a smaller extent than during random foraging (Fig. 1, C and E, and fig. S2). In addition, we observed a new phenomenon that was absent during random foraging. During or preceding runs towards goal locations, some theta sequences swept in the direction of the goal (Fig. 1C and fig. S2A). These goal-directed theta sweeps were intermingled with left and right lateral sweeps (Fig. 1E and fig. S2). Across animals and sessions, we found that 16.3 ± 4.4% (mean ± SD) of theta sweeps in this task were goal-directed (with sweep-to-goal angle ≤ 10º).

Given that animals’ heading direction tended to be correlated with the goal direction in this task, we analyzed the alignment of theta sweeps to heading and goal directions. The percentage of theta sweeps aligned to the goal direction was significantly higher than that to the heading direction (Fig. 1, F and G). The alignment of theta sweeps to the goal direction was not better than the alignment to heading direction during the first trial (*p* = 0.46, bootstrap test) but improved as a function of learning (*p* < 1e-4 from trial 10 to 30, bootstrap test; Fig. 1H). Furthermore, the fraction of goal-directed sweeps gradually increased across learning trials (Fig. 1I), while the fraction of lateral alternating sweeps remained constant (fig. S2D).

As rats could run to any of three rewarded locations, we asked whether theta sweeps predicted which goal location would be visited next. Indeed, theta sweeps showed a high prediction accuracy of the next visited goal, significantly better than using goal-label shuffled data or the prediction from heading direction (Fig. 1J). We specifically analyzed theta sweeps preceding animal choice of trajectory to a specific goal when the heading direction was biased to the goal that would not be chosen. In these cases, goal-directed sweeps were still biased to the chosen goal (Fig. 1, K and L), thus anticipating upcoming spatial trajectories independently of current heading.

Overall, we identified two distinct types of theta sweeps, left-right alternating sweeps and goal-directed sweeps, and showed that their occurrence is modulated by behavioral demands and predicts chosen routes during memory-guided navigation.

### Egocentric goal-direction tuning in CA1 cells and theta sweeps

A mechanism involving inputs from head-direction cells together with spike-frequency adaptation has been proposed to explain left-right alternating theta sweeps (*15, 16*). However, this mechanism cannot explain the goal-directed forward sweeps we observed. While place cells and interneurons have been shown to prominently encode rewarded locations, such a code can only be exploited when an animal is close to the goal (*22, 25–28*) and thus cannot drive goal-directed theta sweeps from locations distant to the goal (Fig. 1, C and K). We hypothesized that an egocentric goal-direction signal could bias theta sweeps towards the goal from any maze location, regardless of heading direction.

To investigate the mechanisms of goal-directed theta sweeps, we first examined the tuning of individual CA1 cells to the goal direction. Rats sampled all goal-direction angles during the goal-directed navigation task (Fig. 2, A and B). Given that most hippocampal cells are tuned to both spatial location and heading direction, we used a linear-nonlinear Poisson (LN) model (*29, 30*) to identify cells whose firing prediction was significantly improved by incorporating goal direction, beyond spatial location and heading direction alone. We found a subset of cells (10.7 ± 3.8%, mean ± SD) that displayed significant angular tuning to the goal direction (Fig. 2, C and D). At the population level, goal-direction tuning spanned all angles, but was more prominent when goal was aligned with heading direction (Fig. 2C). Some goal-direction cells also showed clear place fields, and others exhibited angular tuning to the goal without clear place tuning (Fig. 2E). Although this type of response has not been previously described in rodents, it resembles similar responses observed in flying bats (*31*).

**Fig. 2.**
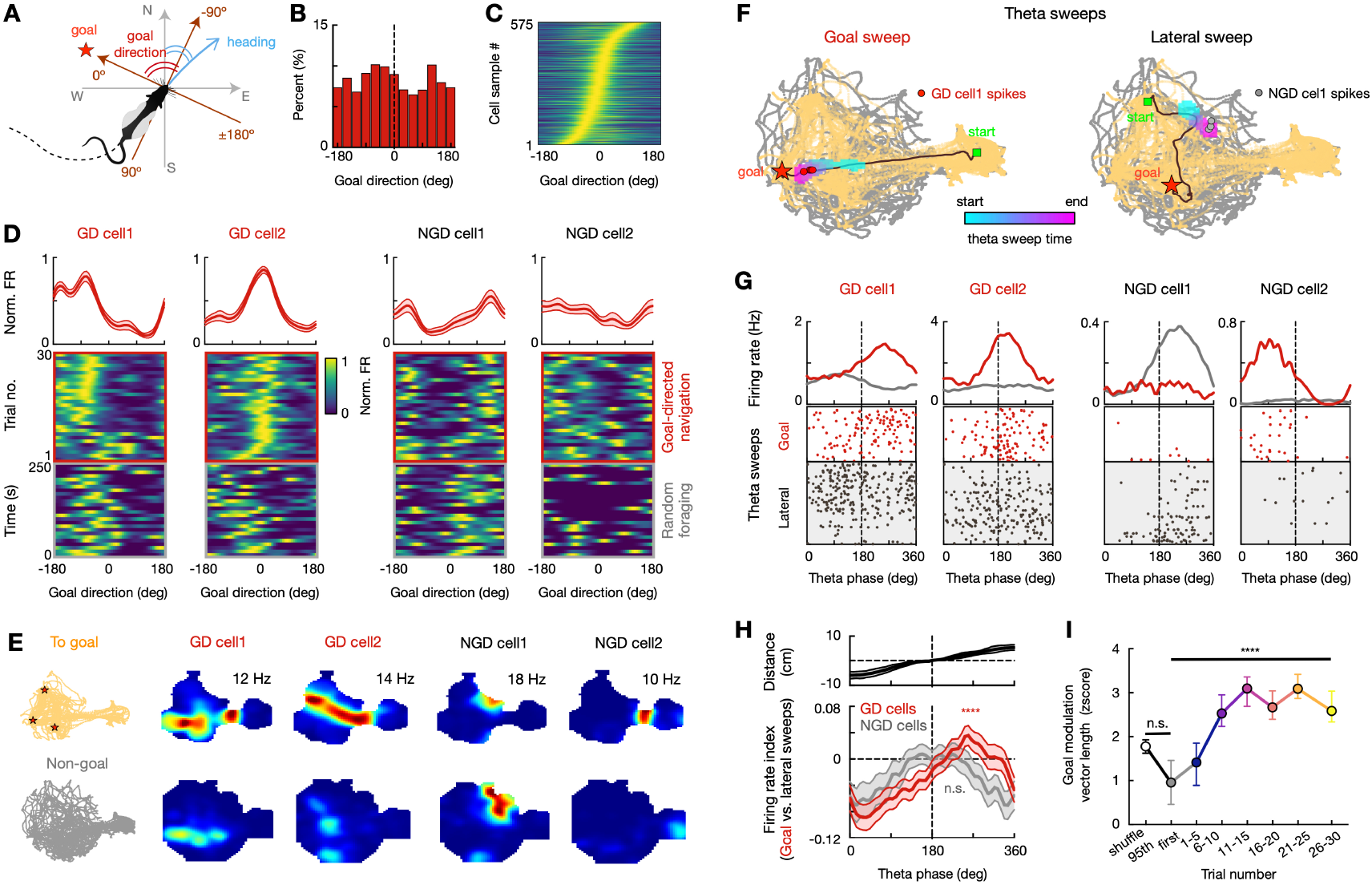
An egocentric goal signal develops over learning during goal-directed theta sweeps. **(A)** Goal-direction angle, defined as the azimuthal angle between the heading direction (blue arrowhead) and the animal-to-goal direction (red arrowhead; (*31*)). **(B)** Distribution of time spent by the animal in different goal-direction angles. **(C)** Normalized goal-direction tuning for all goal-tuned cells sorted by preferred direction (*n* = 575 cell samples from 5 rats; a given cell could contribute up to three cell samples with 3 different goals). **(D)** Four example CA1 cells (*columns*; GD cell: goal-direction cell; NGD cell: non-goal-direction cell). *Top*: Goal-direction tuning curves during *gdn* (mean ± SEM). *Bottom*: Firing rate as a function of goal-direction angles across trials during *gdn* (red square) and during *rf* (below, using goal locations from the *gdn* task). **(E)** Spatial firing rate maps of the cells in **D**. GD cell 1: tuned to both goal direction and place. GD cell 2: tuned to goal direction, weak place tuning. NGD cell 1: classical place cell. NGD cell 2: remaps between *gdn* and *rf*. **(F)** Example goal-directed sweep (*left*) and lateral sweep (*right*) showing spike times (circles) of GD and NGD cells within a theta sweep. **(G)** Theta-phase-aligned rasters and average firing rates of example GD and NGD cells during goal (red) and lateral (gray) sweeps in the same session. **(H)** Firing rate index (goal vs. lateral sweeps) as a function of theta phase (mean ± SEM). *Top*: Distance between decoded and actual position averaged over the theta cycles. *Bottom*, GD cells show elevated late theta phase firing (\*\**p* = 0.0023, *n* = 191 cells from 5 rats), but NGD cells did not (n.s., *p* = 0.46, *n* = 126 cells from 5 rats). Mean firing rates were similar between GD and NGD cells during both goal-directed and lateral sweeps (*p* = 0.50 and 0.61, respectively, rank-sum test). **(I)** Goal-direction modulation (Rayleigh vector length) increased with learning (median ± 95% CI; n.s., *p* > 0.99; \*\*\*\**p* = 1.78e-12; Friedman test with Dunn’s *post hoc*).

Next, we investigated whether the firing of goal-direction cells was coupled with goal-directed theta sweeps. Goal-direction cells fired more during goal-directed compared to lateral sweeps (Fig. 2, F and G). Moreover, goal-direction cells fired more on the late part of the theta cycle during goal-directed sweeps, compared to non-goal-tuned cells (Fig. 2H). The late part of the cycle is when theta sequences sweep ahead towards the goal, while the earlier part of the cycle encodes locations closer to the animal (Fig. 2H). Thus, these observations support a potential role of goal-direction cells in biasing theta sweeps towards goals. Furthermore, the goal-direction tuning of CA1 cells was not significantly different from shuffles in the initial trials but increased rapidly afterwards (Fig. 2I), indicating that goal-direction tuning in CA1 neurons requires learning.

### A circuit mechanism for goal-directed theta sweeps

We next sought to investigate the circuit mechanisms for goal-directed theta sweeps. Previous computational modeling work suggested that spike-frequency adaptation drives the generation of alternating lateral theta sweeps by preferentially inhibiting place cells along the travelled path (*15, 16*). We thus hypothesized that a reduced spike-frequency adaptation together with a goal-direction signal could support the generation of goal-directed theta sweeps. To test this hypothesis, we incorporated variable spike-frequency adaptation and an egocentric goal-direction signal in a continuous attractor network model that can generate alternating lateral theta sweeps (Fig. 3A; (*15*)). With this modification, the model generated both lateral alternating and goal-directed sweeps (Fig. 3B), resembling our experimental data (Fig. 1C). In particular, the switch from lateral to goal-directed sweeps occurred in theta cycles with reduced spike-frequency adaptation and enhanced goal-direction activity during the late half of the cycle (Fig. 3, C and D). These results suggest that a local egocentric goal-direction signal, together with reduced spike-frequency adaptation, could support the generation of goal-directed theta sweeps.

**Fig. 3.**
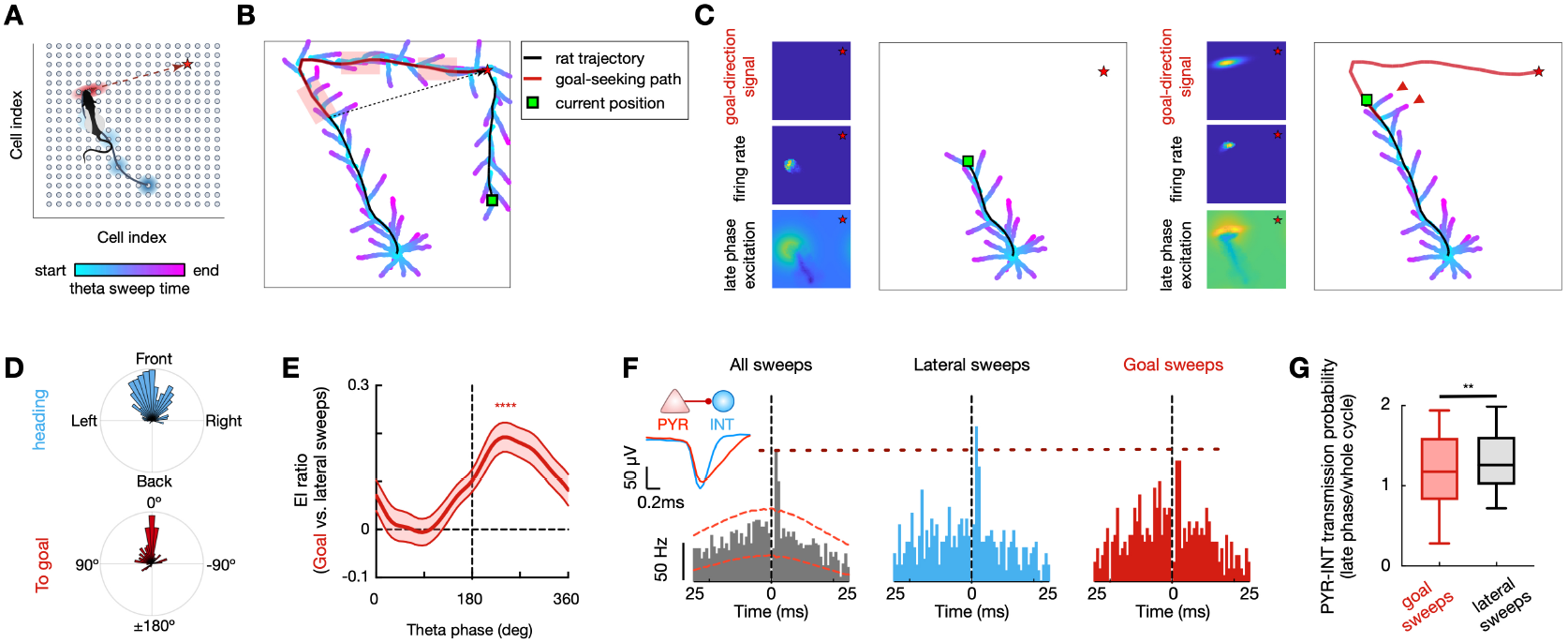
Emergence of goal-directed theta sweeps mediated by reduced feedback inhibition and an egocentric goal signal. **(A–D)** Computational model of goal-directed theta sweeps. **(A)** Schematic of the model, a single-bump, two-dimensional attractor network in which bump movement (blue circle) is driven by spike-frequency adaptation (*15*). Additionally, a subset of neurons exhibits goal-direction tuning (red ellipse), active during late theta phases. **(B)** Simulated theta sweeps, colored by time within each sweep. Goal-direction cells were active during the red segments. Black line: simulated rat trajectory. Red star: goal. **(C)** Example snapshots showing goal signal strength, population firing rate, and excitation strength during the late theta phases before (*left*) and after (*right*) goal signal activation. Red arrowheads highlight goal-directed sweeps. **(E)** Firing rate ratio of CA1 pyramidal cells (PYRs) vs. interneurons (INTs) during goal-directed vs. lateral sweeps, across theta phases (mean ± SEM). Note the increased excitation-inhibition ratio during late theta phases (\*\*\*\**p* = 7.10e-6, rank-sum test, *n* = 36 sessions from 5 rats). **(F)** Cross-correlograms of a representative pre-/postsynaptic PYR–INT pair restricted to the late phase of all theta sweeps, lateral sweeps, or goal-directed sweeps. Red dashed curves: significance bounds. Dashed lines for reference. Inset: averaged spike waveforms. **(G)** Decreased PYR–INT spike transmission probability during late theta phases of goal-directed sweeps compared to lateral sweeps (\*\**p* = 0.0016, *n* = 242 pre-/postsynaptic PYR–INT pairs from 5 rats; Wilcoxon paired test).

We next examined the model predictions in our hippocampal data. A candidate to modulate spike-frequency adaptation in place cells is feedback inhibition from local interneurons (*32, 33*). To determine the role of local inhibition during goal-directed and lateral theta sweeps, we analyzed it as a function of theta phase. Theta sequences sweep from positions behind the animal in the early half of the cycle to positions ahead of it in later theta phases (Fig. 2H, (*5–7*)). Consistent with the model prediction, we found an enhanced excitation/inhibition balance (i.e., reduced inhibition) in the late phase of the theta cycle for goal-directed compared to lateral theta sweeps (Fig. 3E). To specifically examine the contribution of feedback inhibition, we detected functional monosynaptic pyramidal-interneuron (PYR-INT) cell pairs (*34, 35*). PYR-INT spike-transmission probability in the late theta cycle was lower during goal-directed compared to lateral theta sweeps (Fig. 3, F and G), suggesting a reduced feedback inhibition. The concurrent increase in goal-directed cell firing (Fig. 2H) and reduction in feedback inhibition (Fig. 3E–G) during goal-directed, relative to lateral sweeps, supports our model’s prediction of the mechanism underlying goal-directed theta sweep generation.

### Goal-directed theta sweeps were preferentially replayed and coordinated with the prefrontal cortex

During pauses in locomotion, the hippocampus generates temporally compressed place cell sequences that “replay” their ordered firing during theta oscillations in recent behavior (*36–38*). Hippocampal replay has been proposed to support learning and memory formation (*39–45, 18*), and its generation is linked to plasticity during theta sequences (*13, 46*). What determines that specific aspects of experience (e.g., certain routes) are prioritized during replay is not understood (*13, 47–52*). We hypothesized that, if goal-directed theta sweeps play a role in memory-guided navigation, they might be preferentially replayed compared to lateral sweeps or other experienced spatial trajectories. To investigate the relationship between theta sweeps and replay, we detected sequential activity patterns during immobility periods in the maze that coincided with sharp-wave ripples (SWRs), and compared the spatial trajectories encoded during those replay events with those encoded during theta sweeps. Consistent with our hypothesis, replay sequences were preferentially biased towards goal locations (Fig. 4, A and B, and fig. S3), mirroring the structure of goal-directed theta sweeps (Fig. 4C). Moreover, replay sequences were significantly more similar to goal-directed theta sweeps than to lateral sweeps (Fig. 4D), suggesting that replay preferentially reinstates goal-directed theta sweeps.

**Fig. 4.**
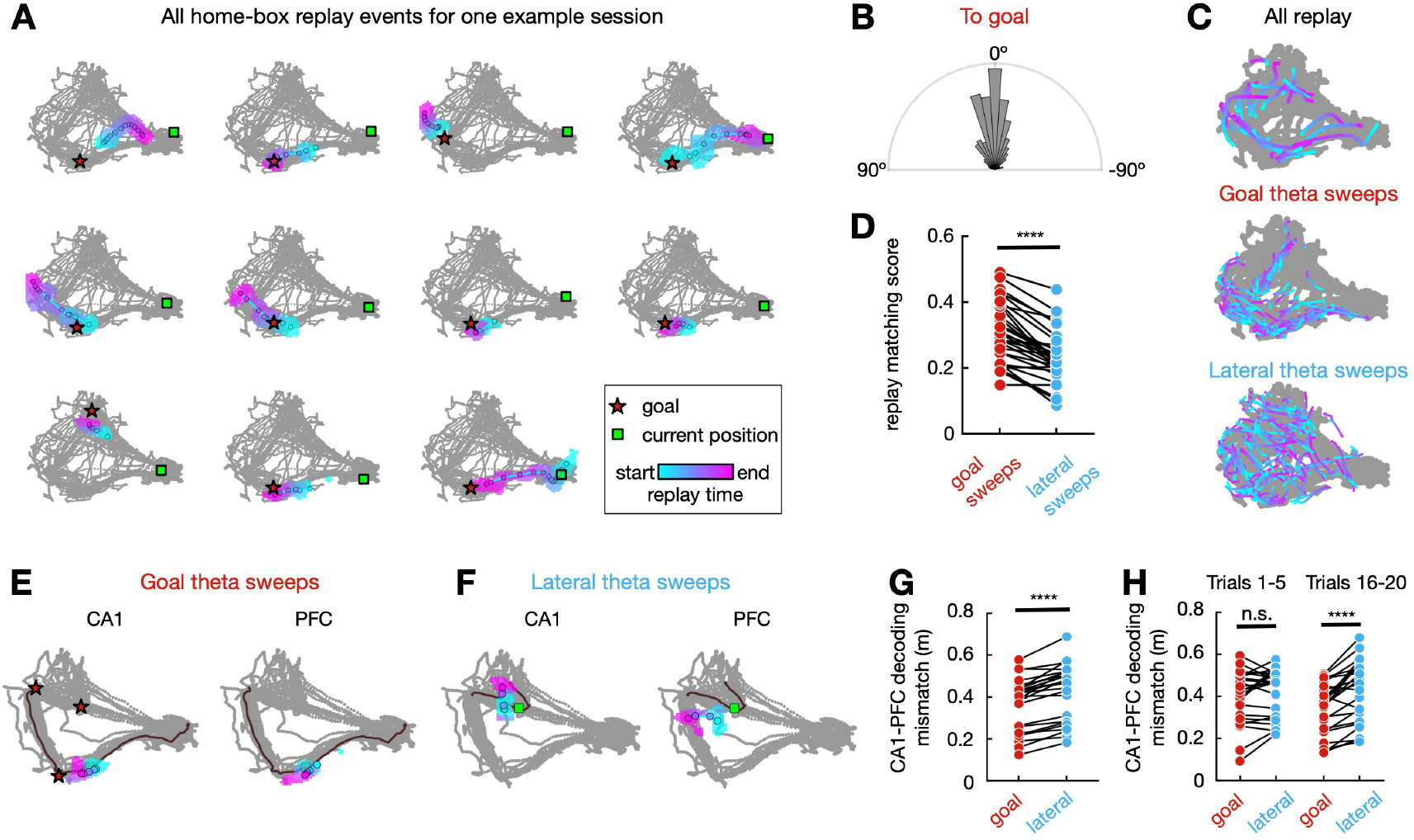
Goal-directed theta sweeps are preferentially replayed and coordinated with the prefrontal cortex. **(A)** All significant replay events (*n* = 11) occurring in the home box during a representative *gdn* session. Colored blobs represent decoded positions during replay events (filled circles: positions with maximal decoded probability). Note that most replay events were directed toward goal locations (red stars: closer goal to replay decoded trajectory; green square: current animal position, gray lines: animal trajectories in the whole session). **(B)** Polar distribution of replay direction relative to goals during *gdn* (*n =* 476 replay sweeps from 5 rats). Plots span half a cycle, as forward and reverse replay directions were collapsed using absolute angles. **(C)** All awake replay events, goal-directed theta sweeps, and lateral sweeps from the example session shown in **A** (each color-coded line for one event). **(D)** Decoded positions during replay were more strongly correlated with those during goal-directed than lateral theta sweeps (\*\*\*\**p* = 4.66e-9, *n* = 31 sessions from 5 rats, Wilcoxon paired test). **(E-F)** Example goal-directed **(E)** and lateral **(F)** CA1 (*left*) and PFC (*right*) theta sweeps showing decoded trajectories (similar presentation as in **A**). **(G)** Greater mismatch between CA1 and PFC decoded positions during goal-directed sweeps compared to lateral sweeps (\*\*\*\**p* = 2.38e-7, *n* = 24 sessions in 4 rats, Wilcoxon paired test). **(H)** CA1-PFC coordination during goal-directed theta sweeps developed over learning (n.s., *p* = 0.30 for trials 1-5, and \*\*\*\**p* = 1.05e-5 for trials 16-20, Wilcoxon paired test).

If goal-directed theta sweeps support planning of upcoming spatial trajectories, we would expect them to be coordinated with other brain regions also involved in planning and decision making, such as the PFC. PFC ensemble activity in theta cycles encoded compressed spatial trajectories sweeping ahead of the animal in the goal-directed task (Fig. 4, E and F, and fig. S3B), similar to hippocampal theta sweeps. PFC theta sweeps were more coordinated with the hippocampus, i.e. represented more similar spatial trajectories, during goal-directed CA1 sweeps than during lateral sweeps (Fig. 4E-G and fig. S3). Furthermore, the coordination of PFC sweeps with hippocampal goal-directed sweeps increased with learning (Fig. 4H).

### Theta sweeps exploit internal latent maps of the environment

In line with the traditional view that the hippocampus predominantly encodes physical space (*1, 2*), theta sequences could be a mechanism for rapid sampling of the surrounding environment (*14*). An alternative possibility is that the hippocampus encodes different maps of the same environment that correspond to different behavioral contexts, and theta sequences are a mechanism to explore these internal representations and guide flexible behavior. According to the first hypothesis, if an animal learns two different sets of rewarded locations in the same maze, its hippocampus may preserve the same global map in both conditions, albeit with some degree of place cell changes around rewarded locations (Fig. 5A) (*22, 27, 28, 53*). Alternatively, the hippocampus may generate two different maps representing each reward configuration (or “latent” maps, since environmental features do not change; Fig. 5A). In the second case, theta sequences would be embedded within these latent representations and dynamically switch as behavioral context changes, rather than traversing a common spatial map in both cases.

**Fig. 5.**
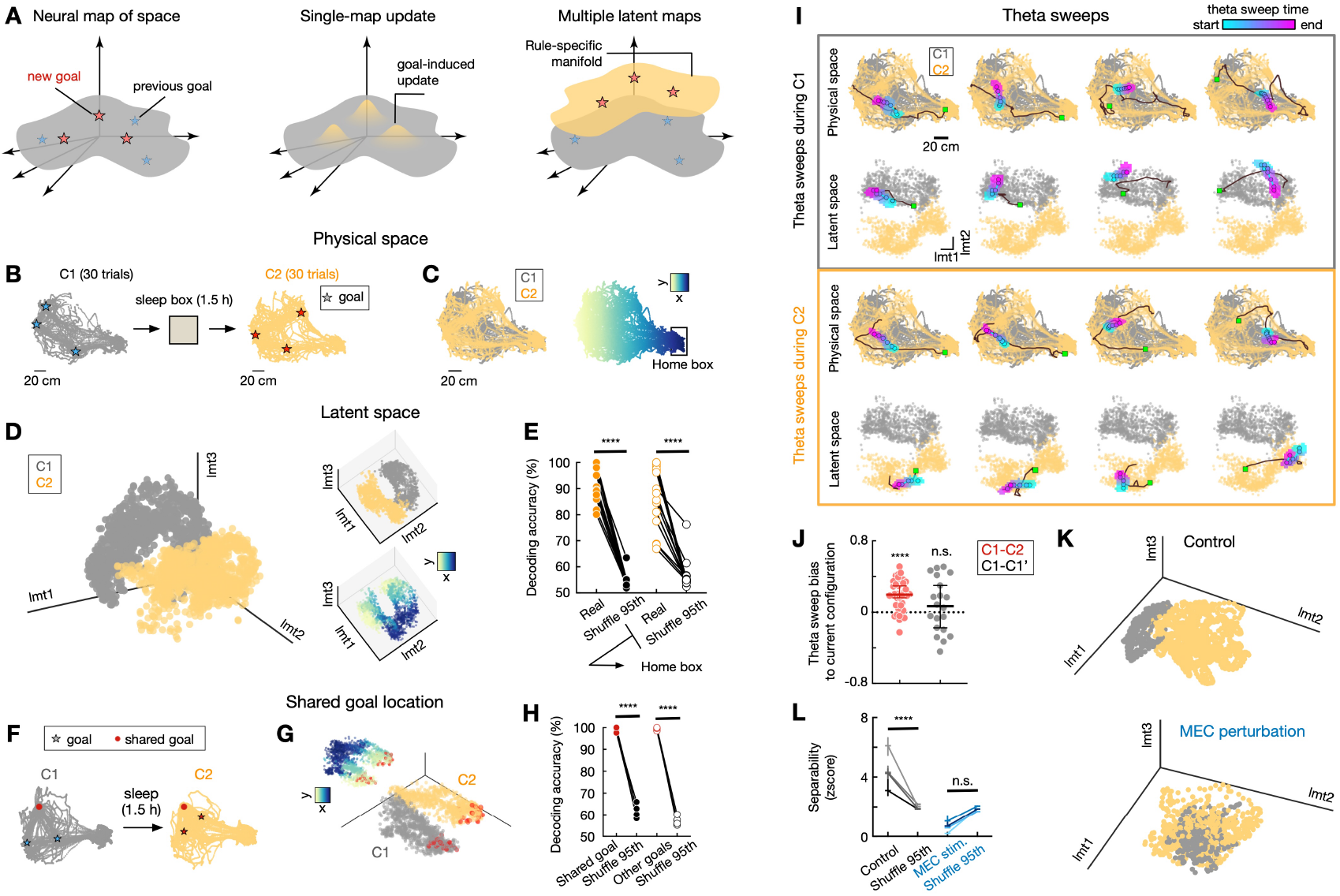
Theta sequences exploit distinct context-specific latent maps in the hippocampus. **(A)** When learning a new set of hidden reward locations (goals), the hippocampus may either update an existing map (*middle*, “single-map update”) or form distinct, coexisting latent maps for each condition that are selectively engaged based on context (*right*, “multiple latent maps”). **(B)** Experimental design: animals learned configuration 1 (C1), followed by 1.5-hour rest, then configuration 2 (C2). **(C)** Animal trajectories, color-coded by configuration (*left*; C1, gray; C2, orange) and by position (*right*). **(D)** Three-dimensional embedding of neural activity using LMT, colored by configuration and position (*inset*). Two distinct manifolds correspond to the two configurations. **(E)** Condition decoded from neural activity using *k*-nearest neighbor (kNN) and 10-fold cross-validation. Decoding accuracy exceeded that of shuffled session labels (\*\*\*\**p* = 1e-4), even when decoding from home box activity alone (\*\*\*\**p* = 8.78e-9; *n* = 19 C1–C2 session pairs from 5 rats; one-way repeated-measures ANOVA). **(F–H)** Shared goals were represented in distinct latent states across conditions. **(F)** Task design with a shared goal between C1 and C2. **(G)** Manifold embedding showing separate latent states for visits to the shared goal (red circles). *Inset*: same embedding, color-coded by position. **(H)** Task configuration decoding from neural activity at the shared goal location and other goals (\*\*\*\**p* = 6.00e-5 for shared goal, 6.54e-6 for other goals; *n* = 5 session pairs from 3 rats; one-way repeated-measures ANOVA). **(I)** Example theta sweeps decoded using standard spatial rate maps (*top rows*) and LMT-based latent tuning curves (*bottom rows*). Sweeps traversed similar spatial paths but exploited configuration-specific latent maps. **(J)** Theta sweep decoding was strongly biased toward the current configuration (\*\*\*\**p* = 4.63e-7; *n* = 38 sessions from 5 rats) and was not explained by temporal drift (C1’, re-exposure to C1: n.s., *p* = 0.33; *n* = 20 sessions from 4 rats; rank-sum test vs. 0). **(K–L)** Optogenetic disruption of MEC spike timing, which impaired theta sequences (*13*), also impaired latent map formation. **(K)** Manifold embeddings from an example control and MEC perturbation sessions. **(L)** Separability of condition-specific latent maps, measured by Dunn’s index, was reduced during MEC perturbation (all *p’s* > 0.05 for MEC perturbation sessions, and all *p’s* < 9.3e-14 for control sessions compared to context-label shuffles, Wilcoxon paired test; *n* = 4 sessions from 3 rats per condition).

To distinguish between these two possibilities, we trained rats to perform the same goal-directed navigation task as before but with two different reward configurations in consecutive sessions on the same day (Fig. 5B). Place cell tuning remained largely stable across the two sessions (fig. S4). Some place fields near old and new rewarded locations changed their firing rates or shifted towards newly rewarded locations on the second session (fig. S4), as previously described (*22, 23*). Overall, the hippocampal spatial representation was largely preserved across task configurations (fig. S4).

To distinguish between the “single-map updating” and “multiple latent maps” hypotheses (Fig. 5A), we analyzed the representational structure of ensemble neural activity, or neural manifold, across task configurations. We applied Latent Manifold Tuning (LMT; (*54*)), an unsupervised method that simultaneously estimates neural manifolds and neuronal tuning to latent variables. If the hippocampus encodes multiple latent maps of the same physical maze, then distinct reward configurations should be separable based on population activity, even at identical physical locations throughout the maze, not just near the goals (Fig. 5A). Consistent with this prediction, we found that, despite only modest changes in single-cell place fields across sessions, population-level dynamics revealed distinct latent maps for each reward configuration within the same maze (Fig. 5D). Importantly, the same physical locations were represented as different latent states depending on the task context, including in regions far from the goals, such as the home box (Fig. 5D). As a result, the current reward configuration could be accurately decoded from neural activity (Fig. 5E). Furthermore, even when the two reward configurations shared a common goal location (Fig. 5F), neural activity at the shared goal diverged across the two latent maps (Fig. 5, G and H), indicating a distinct context-dependent coding of the same goal.

Place cell responses in the same environment gradually change over time, or drift, even in the absence of external changes (*29, 55, 56*). To test whether such temporal drift can explain the generation of distinct maps of the same maze, we re-exposed rats to the second reward configuration in a new session after 1.5 hours of rest. The distance of the neural subspace in the second task session with the previous one (with a different reward configuration) was larger than compared with the re-exposure session (with the same reward configuration) (fig. S5). This indicates that temporal drift alone cannot explain the generation of distinct maps for the different reward configurations.

Next, we examined the dynamics of theta sweeps across sessions with different reward configurations. According to the latent-map hypothesis, we would expect theta sequences to sweep only within the relevant latent map for each configuration. To test this, we decoded theta sweeps in the latent space comprising the subspaces of both task configurations. Theta sweeps from one of the configurations were restricted to the corresponding neural subspace for that configuration (Fig. 5, I and J). Even for the theta sequences sweeping along similar spatial trajectories in both sessions, they were completely segregated in neural space (Fig. 5I). In a subset of sessions, animals encountered the same reward configuration across two sessions. For those sessions, theta sweeps did not show a bias to the subspace of the current session (Fig. 5J and fig. S6), suggesting that the separation of theta sweeps for different task configurations could not be simply explained by temporal drift. Overall, these results suggest that theta sweeps are a mechanism to explore internal representations of task space.

Finally, we analyzed previously published data in which we disrupted CA1 theta sequences by optogenetically perturbing the temporal structure of medial entorhinal cortex (MEC) activity while rats performed the same goal-directed cheeseboard maze task (*13*). Disrupting theta sequences during learning trials prevented the generation of two distinct maps for different reward configurations (Fig. 5, K and L). This suggests that synaptic plasticity during theta sequences may be a driving mechanism for the generation of latent spatial representations in the hippocampus.

## Discussion

We found that the hippocampus generates two distinct types of temporally compressed place cell sequences within theta cycles during navigation. In rats randomly foraging in an open maze, theta sweeps alternated left and right of the animal’s location in a highly stereotyped pattern. During goal-directed navigation to previously learned hidden goals on the same maze, some theta sequences swept instead towards one of the goal locations. These goal-directed sweeps were intermingled with lateral sweeps and were independent of heading direction. In contrast to the rigid nature of lateral theta sweeps, goal-directed sweeps were modulated by learning and predicted upcoming spatial trajectories to chosen goals. Goal-directed sweeps can therefore provide a useful vectorial signal to plan routes to goals, even in the absence of local landmarks. This type of signal provides essential complementary information for flexible navigation in addition to other types of codes implemented by different cell types in the entorhinal-hippocampal network (*1, 2, 17, 27, 30, 33, 35, 57–59*).

We propose that both types of theta sweeps support complementary functions during navigation. Lateral sweeps might be a mechanism for unbiased sampling of the surrounding space, including unvisited locations (*14*), and contribute to the generation and updating of internal maps. On the other hand, goal-directed sweeps may facilitate memory-guided navigation and trajectory planning. In support of this interpretation, goal-directed hippocampal sweeps were more coordinated with trajectory representations in the PFC, another region implicated in behavioral planning and goal-directed navigation (*17–21*), than lateral theta sweeps. These results provide a unified framework to explain previous findings, including exclusive lateral sweeps during open field and linear maze tasks (*14*), sweeps to different arms at choice points in linear mazes (*8, 9, 11*) and forward sweeps to reward ports along linear tracks (*10, 12*).

A subset of hippocampal cells developed goal-direction tuning selectivity over learning. These goal-direction cells provided a vectorial egocentric signal pointing to goals, even from distant locations in the maze. They thus provide a complementary mechanism for goal coding to the local goal overrepresentation by place cells commonly observed in rodents (*22, 26–28*), and are reminiscent of similar goal-vector signals found in flying bats (*31*). Computational simulations indicated that a combination of variable spike frequency adaptation (*15, 16*) and an egocentric goal-direction signal in a continuous attractor network model can generate both lateral and goal-directed theta sweeps. In support of the model predictions, we found increased goal-direction cell activity and decreased feedback inhibition to place cells (a mechanism for spike frequency adaptation-(*32*)) during goal-directed theta sweeps.

Hippocampal place cells replay experienced spatial trajectories during pauses in locomotion and sleep, coordinated by SWRs (*13, 36–39, 60*). In our goal-directed navigation task, we found that replay events during immobility periods on the maze were strongly biased towards goal locations and more correlated with goal-directed than lateral theta sweeps. This observation suggests that goal-directed theta sweeps could be a mechanism for promoting the selection of behaviorally relevant spatial trajectories during replay and subsequent memory consolidation (*44, 47–51*).

When rats learned two different sets of rewarded locations on the same maze, the hippocampus generated two distinct representations, or maps, of the maze. Despite modest changes in individual place cell tuning, at the population level, low-dimensional representations of the same maze for both reward configurations were completely segregated. This result is in line with recent evidence suggesting that the hippocampal cognitive map not only reflects physical space and associated sensory cues but also internal variables such as behavioral relevance, expectations or learned rules (*29, 61–63*). Furthermore, theta sweeps along similar spatial trajectories during sessions with different reward configurations were restricted to the corresponding neural subspace. Therefore, theta sweeps could provide a mechanism for mental exploration of internal representation of the world in support of planning, simulation and prediction of flexible behavior.

## Acknowledgments

The authors thank Can Liu for help with surgical procedures and members of the Oliva and Fernandez-Ruiz labs for providing useful feedback on the manuscript. This work was supported by NIH grant R01MH130367 and Whitehall Foundation (AO), NIH grant R01MH136355, DP2MH136496, Sloan Fellowship, Whitehall Research Grant, Klingenstein-Simons Fellowship and Pershing Square Foundation’s MIND Prize (AFR), and Klarman Fellowship (WT).

## Author contributions

W.T, X.M., R.E.H, E.C-L and T.N performed the experiments. W.T, X.M., R.E.H, H.C. and A.F.R. analyzed the data. A.F.R, W.T. and A.O. wrote the manuscript with input from all authors.

A.F.R. and A.O. supervised the work.

