## Supplementary material for "Goal-directed hippocampal theta sweeps during memory-guided navigation": Methods and Supplemental figures

**The PDF file includes:**

Materials and Methods
Figs. S1 to S6
References (*64*-*68*)

**Other Supplementary Materials for this manuscript include the following:**

Movie S1

### Materials and Methods

Animals

All experiments conformed to guidelines established by the National Institutes of Health and have been approved by Cornell University's Institutional Animal Care and Use Committee. Rats ($n = 7$; adult male Long-Evans, 300-500 g, 3-6 months old) were kept in the vivarium on a 12-hour light/ dark cycle. Temperature and humidity in the room were kept at 68-72 F and 40-60%, respectively. They were housed 2 per cage before surgery and individually after it.

Surgical procedures

Silicon probes (NeuroNexus, Cambridge Neurotech, or Diagnostic Biosignals) were mounted on custom-made 3D-printed micro-drives to allow precise adjustment of the vertical position of sites after implantation. The probes were inserted above the target region. Craniotomies were sealed with sterile wax. Two stainless steel screws were placed bilaterally over the cerebellum to serve as ground and reference electrodes. Several additional screws were driven into the skull and covered with dental cement to strengthen the implant. Finally, a copper mesh mounted on a 3D-printed resin base was attached to the skull with dental cement and connected to the ground screw to act as a Faraday cage, attenuating the contamination of the recordings by environmental electric noise and protecting the headgear. After post-surgery recovery, probes were moved gradually in 50 to 150 µm steps per day until the desired position was reached. Hippocampal and cortical layers were identified physiologically by unit activity and characteristic LFP patterns (*19*, *24*, *64*). A variety of different silicon probes were implanted targeting dorsal hippocampal CA1 (-4.0-4.5 mm antero-posterior from Bregma, AP; ± 2.6 mm from midline, ML; 5 rats, right hemisphere; 2 rats, bilateral) and the right PFC (-3 mm AP and 0.7 mm ML; 4 out of 7 rats). For optogenetic experiments (*13*), rats were also implanted with custom-made optic fiber arrays (three 200-µm core multimode fiber each, ~500 µm apart, connected to a single 2.5 mm steel ferrule; Doric Lenses) in both hemispheres over the medial entorhinal cortex (–7.7 / –8.4 / –9.1 AP; ± 4.6 ML and 4.7/ 4.3/3.2 mm from the surface of the brain, for each fiber respectively; see also *optogenetic manipulations*).

Behavioral tasks

After surgery, animals were handled daily and accommodated to the experimenter, recording room, and cables for one week before the start of the experiments. Prior to the start of the behavioral experiment, the animals were water-restricted. Recordings were conducted using the Intan RHD2000 interface board or Intan Recording Controller, sampled at 20 kHz. Amplification and digitization were done on the head stage. In all the behavioral sessions, animals' position was tracked with an overhead camera (Basler) at 30 Hz (Pylon 5.0 software), and later extracted using DeepLabCut (DLC, v2.3.0) (*65*).

The cheeseboard maze was a circular platform (120-cm diameter), where the animals learned to find three goal wells that contained water rewards ($n = 5$ rats). A trial was completed once the animal had retrieved all rewards and returned to the start box to collect an additional food pellet reward. The locations of the goal wells changed daily but were fixed within a session. This strategy required the animals to update their memory for the new goal locations in each session but in an otherwise familiar environment. Note that there was always a delay of approximately 30 seconds between trials. If the rat could not find all three rewards in 2 minutes, the trial was ended, and the animal was placed back in the start box. Each experimental day began with a pre-probe session consisting of five trials using the same reward configuration as the previous day, to assess whether the animal remembered the previous day's reward locations. This was followed by a 1.5-2 hour sleep session. Subsequently, the animal underwent a learning session to acquire a new set of three reward locations (configuration 1, C1), conducted either with or without light stimulation. After a post-learning 1.5-2 hour sleep session, the animal underwent training involving either the acquisition of a novel reward configuration (configuration 2, C2) or the recall of a previously learned configuration (re-exposure to configuration 1, C1'). Each training session consisted of 20 to 30 trials. Learning performance was evaluated as the latency and the distance traveled from the start box to collect the three rewards. For the optogenetic experiments (Fig. 5, K-L, see also *optogenetic manipulations*), stimulation (Stim ON) and control (Stim OFF) sessions were alternated in a counterbalanced design across animals. To identify approach trajectories toward a specific goal, we manually annotated the start and end points of these trajectories based on the animals' tracked positions in 37 out of 63 recorded sessions. In a subset of sessions, animals foraged for randomly baited water rewards on the same cheeseboard, either for 20 minutes following a 1.5-2 hour sleep session, or for 5-10 minutes immediately after the 20–30 trial learning session.

Optogenetic manipulations

For optogenetic experiments, rats were injected with custom-prepared AAV5-mDlx-hChR2(H134R)-mCherry from AddGene (plasmids were a gift from Dr. Gord Fishell; (*66*). Three injections per hemisphere were performed along the dorso-ventral MEC axes as follows: 1) -7.7 AP, ±4.6 ML, 4.7 mm depth, 200 nL; 2) -8.4 AP, ±4.6 ML, 4.3 mm depth, 400 nL; 3) -9.1 AP, ±4.6 ML, 3.2 depth, 700 nL. After injection, craniotomies were sealed and animals recovered in the vivarium for three weeks. Following this period, a second surgical procedure for implanting optic fibers and electrodes was performed, as described above. Optic fiber arrays were implanted in the same craniotomies performed previously for virus injection. For optogenetic stimulation, fiber array ferrules were connected with mating sleeves to 450-nm blue light-emitting laser diodes coupled to 2.5 mm steel ferrules (PL-450, Osram).

Optogenetic perturbations were performed by delivering blue light, modulated with a positive 53 Hz current sinusoid using an isolated current driver (Thorlabs). Light intensity was calibrated for each animal during home cage recordings by analyzing the suppression of LFP gamma power in the *stratum lacunosum-moleculare* during stimulation. A minimum power of 3 mW and a maximum of 6 mW was used. In the goal-directed navigation task on the cheeseboard, light stimulation was applied during all running periods in the maze during learning trials.

Recording system and data preprocessing

An Intan RHD2000 interface board or Intan Recording Controller was used for electrophysiological recordings. The sampling rate was set at 20 kHz. Both amplification and digitization were done in the head stage (Intan Technologies). Data were visualized online during recording using the Intan software and Neuroscope (Neurosuite). LFP signals were down-sampled at 1250 Hz for subsequent analysis.

Tissue processing and immunohistochemistry

Following the termination of the experiments, animals were deeply anesthetized and perfused transcardially first with 0.9% saline solution followed by 4% paraformaldehyde solution. The brains were sectioned into 70-μm thick slices (Leica Vibratome). The sections were washed and mounted on glass slides with a fluorescence medium (Fluoroshield with DAPI - F6057, Sigma, USA). A confocal microscope (Zeiss LSM 800) was used to obtain high-quality photos.

### Quantification and statistical analyses

#### Spike sorting and single unit classification

Spike sorting was performed semi-automatically using KiloSort (https://github.com/cortex-lab/KiloSort), followed by manual curation using the software Phy (https://github.com/kwikteam/phy) and custom designed plugins (https://github.com/petersenpeter/phy-plugins) to obtain well-isolated single units. Cluster quality was assessed by manual inspection of waveforms and auto-correlograms, and by isolation distance metrics. Multi-units, noise clusters, or poorly isolated units were discarded from further analysis. Well-isolated units were classified into putative cell types using the MATLAB package, Cell Explorer (*67*) (https://github.com/petersenpeter/CellExplorer). Spiking features such as auto-correlogram (ACG), spike waveform, and putative monosynaptic connections derived from short-term cross-correlograms (CCGs), were used to characterize and classify well-isolated units (*19*, *43*). Three cell types were assigned: putative pyramidal cells, narrow waveform interneurons, and wide waveform interneurons (fig. S1). The two key metrics used for this separation were burst index and trough-to-peak latency. Burst index was determined by calculating the average number of spikes in the 3-5 ms bins of the spike ACG divided by the average number of spikes in the 200-300 ms bins. To calculate the trough-to-peak latency, the average waveforms were taken from the recording site with the maximum amplitude for the averaged waveforms of a given unit. Only putative pyramidal cells were used for further analysis, unless otherwise specified.

#### Spatial fields and linearization

Spatial fields were calculated only during running periods (> 5 cm/s) at positions with sufficient occupancy (> 20 ms). 2D occupancy-normalized rate maps were constructed using spike counts and occupancies with 50 x 50 spatial bins, smoothed with a 2D Gaussian kernel (SD = 2).

#### Population vector correlation

To measure remapping in population activity (fig. S4C), a population vector (PV) was constructed as the firing rate vector of all spatial-tuned cells in a certain spatial bin. The PV correlation was then defined as the Pearson correlation ($r$) between the PVs across all bins in the mazes between two sessions.

#### Theta phase estimation

To extract the theta phase, one channel around the CA1 *stratum oriens* was chosen and band-pass filtered in the range of 5-15 Hz. The theta phase was then computed using the Hilbert transform of the filtered LFP. Since the theta phase can vary depending on the exact position of the channel within the hippocampus, theta phase was then re-aligned such that the distance between the decoded position (see *Decoding theta and replay sweeps with Bayesian reconstruction*) and the animal position was maximal at zero phase (Fig. 2H; (*14*, *15*)).

#### SWR and HSE detection

For SWR detection, the wide-band signal from a CA1 pyramidal layer channel was filtered (difference of Gaussians; zero-lag, linear-phase finite impulse response (FIR) filter), and instantaneous power was calculated by clipping at 4 SD, rectified and low-pass filtered (*13*, *43*, *44*, *64*). The low-pass filter cut-off was at a frequency corresponding to π cycles of the mean band-pass (for 80–250 Hz band pass, the low-pass was 55 Hz). Subsequently, the power of the non-clipped signal was computed, and all events exceeding 4 SD from the mean were detected. The events were then expanded until the non-clipped power fell below 1 SD sharp waves were detected separately using LFP from a CA1 *stratum radiatum* channel, filtered with band-pass filter boundaries (5–40 Hz). LFP events of a minimum duration of 20 ms and a maximum of 400 ms exceeding 2.5 SD of the background signal were included as SWRs.

To refine the detection of SWR start and end for replay analyses (see *decoding theta and replay sweeps with Bayesian reconstruction*), the high synchrony events (HSEs) during immobility period (less than 5 cm/s) were first detected as previously reported (*13*, *44*). To detect HSEs, the combined activity of all pyramidal spikes was binned into 1-ms bins and smoothed with a Gaussian kernel (SD = 15 ms) to produce a population firing rate curve over time. HSEs were initially detected as contiguous periods when the population firing rate stayed above 3 SD of the mean, and further refined as times around the initially detected events during which the firing rate exceeded the mean. Only HSEs with a duration of 50 ms or more were included for further replay analysis.

**Classification of goal-direction cells**

To detect the goal-directional cells, we first compute the goal direction ($\gamma$) (*31*), defined as the angle between the azimuth heading direction of the rat ($\varphi$) and the direction of the vector ($\alpha$) connecting the head position ($x$, $y$) with the goal position ($x_G$, $y_G$) as follow:

$$\alpha = angle\big((x_G - x) + (y_G - y) * i\big),$$
$$\gamma = modulo\big((\alpha - \varphi) + \pi, 2\pi\big) - \pi.$$

The goal-direction tuning curves of each cell were then computed by counting the number of spikes in each azimuthal bin (51 bins) and dividing it by the total time spent in that bin during all running periods in the maze during learning trials and smoothed with a Gaussian kernel (SD = 1.5 bins) in a circular manner. The goal modulation was then measured as the Rayleigh vector length based on the goal-direction tuning curves, and the significance of goal-direction tuning was assessed by circularly time-shifting all spikes by a random interval, wrapping the end of the session to the beginning (*31*).

To further rule out potential confounds from place ($p$) and egocentric heading ($\varphi$) signals in the identification of goal-direction cells we used the LN model (*30*) to estimate the spike rate of a neuron at time bin $t$ ($r_t$) as an exponential function of the sum of all variables ($p_t$, $\varphi_t$, $\alpha_t$) projected onto a corresponding set of parameters ($\beta_p$, $\beta_\varphi$, $\beta_\alpha$) as:

$$\boldsymbol{r} = exp(\textstyle\sum_i X_i^N \beta_i)/\Delta t,$$

where $\boldsymbol{r}$ denotes a vector of firing rates of a neuron across $N$ time bins, $i$ indexes the variables ($i \in [p, \varphi, \alpha]$), and $X_i^N$ is a state matrix for the $i$-th variable. Each column of $X_i^N$ is a state vector $x_i$ at each of the $N$ time bins, whose elements are 0, except for the element (= 1) corresponding to the animal's current state of that variable.

To learn the parameters ($\beta_p$, $\beta_\varphi$, $\beta_\alpha$), we maximized the Poisson log-likelihood of the observed spike train given the model-estimated spike number $r\Delta t$. The parameters were optimized using the *fminunc* function from MATLAB, and the log-likelihoods and variance explained were calculated from the held-out data via 10-fold cross-validation. To select the best LN model that characterizes the single-neuron response, we compared both the log-likelihoods and variances explained of the reduced ($p$, $\varphi$) and the full model ($p$, $\varphi$, $\alpha$). Statistical significance was assessed using one-sided rank-sum tests ($p < 0.05$) (*30*). A neuron was classified as goal-directional if the full model showed significantly higher log-likelihood and explained variance compared to the reduced model (i.e., both $p$'s $< 0.05$). Neurons with non-significant differences on both measures (both $p$'s $> 0.05$) were classified as non-goal-directional (Fig. 2).

**Decoding theta and replay sweeps with Bayesian reconstruction**

We used a previously established method for detecting left-right lateral theta sweeps with Bayesian decoder to detect theta sweeps in our dataset (*14*). In brief, the spike times of CA1 pyramidal cells were binned into 10-ms bins, and the position was decoded in each time bin from the matrix of firing rate and tuning curves from $N$ neurons, with an assumption of Poisson firing and a uniform prior:

$$P(\boldsymbol{x}|\boldsymbol{y}) \propto exp\left(\sum_{i=1}^{N} y_i log\left(f_i(\boldsymbol{x})\right) - dt \sum_{i=1}^{N} f_i(\boldsymbol{x})\right),$$

where $P(\boldsymbol{x}|\boldsymbol{y})$ is the conditional probability for the rat's 2D location $\boldsymbol{x}$, given the observed spike count $\boldsymbol{y}$ and tuning curve $f(\boldsymbol{x})$. The decoded position was taken as the position bin that maximized $P(\boldsymbol{x}|\boldsymbol{y})$.

We then extracted individual sweeps from the decoded position trajectory by a simple sequence detection algorithm, as previously reported (*14*). Within each theta cycle, candidate sweeps were identified as the longest sequence of consecutive valid 10 ms time bins in which the decoded position shifted by less than 20 cm and the direction changed by less than 90° between adjacent bins. These candidate sweeps were then truncated to maximize the net Euclidean distance from start to end. A sweep vector $\boldsymbol{s}$ was defined as the vector connecting the low-pass filtered decoded position at the start of the theta cycle to the most distal point of the candidate sweep. To assess whether the decoded trajectory followed a coherent and straight path, we computed the goodness-of-fit ($r^2$) between the sweep vector axis and the series of $[x, y]$ coordinates comprising the sweep, as previously described (*14*). Candidate sweeps were retained for further analysis if they consisted of at least four samples and exhibited a goodness-of-fit ($\boldsymbol{r^2}$) greater than 0.5. Sweep direction and length were defined as the angle and magnitude, respectively, of the corresponding sweep vector. Sweep prevalence was calculated as the proportion of theta cycles, during which the animal's running speed exceeded 15 cm/s, that contained sweeps meeting these criteria. Replay sweeps were extracted using the same method as for theta sweeps, but during HSEs coinciding with SWRs with running speed < 5 cm/s (see *SWR and HSE detection*).

**Goal prediction with theta sweeps**

We used a multi-class SVM decoder ($k = 3$ goals) to predict the animal's choice of goals on the current path or trial, based on the sweep direction relative to either the goals or the heading direction (Fig. 1J). For each theta sweep, we computed the sweep vector ($s$), heading vector ($h$), and goal vectors ($\alpha$), defined as vectors connecting the animal's head position ($x$, $y$) to each of the three goal positions ($x_G$, $y_G$). The sweep-to-goal angle was defined as the angle between the sweep vector ($s$) and a goal vector ($\alpha$), while the sweep-to-heading angle was defined as the angle between the sweep vector ($s$) and the heading vector ($h$). We included all theta sweeps detected along goal-approaching paths (Fig. 1A) and trained two separate multi-class SVM decoders using MATLAB's *fitcecoc* function: one using sweep direction relative to the goals and another using sweep direction relative to the heading. Each decoder aimed to predict the upcoming goal from three possible options. Decoding accuracy was evaluated using 5-fold cross-validation on held-out test data. To assess chance-level performance, we created shuffled datasets by randomly permuting goal identities across goal-approaching paths.

**The LMT model**

Using the spatial firing rate maps, we can only decode position in one environment based on the tuning curves referenced to the physical space, making it impossible to detect context-specific representations and theta sweeps. To simultaneously extract the neural representations for different behavioral contexts in the same maze and characterize the neuronal tunings to latent task variables, we therefore adapted the latent manifold tuning (LMT) method (*14*, *54*). In this framework, all the simultaneously recorded CA1 pyramidal cells that pass the cell inclusion criteria (see *spike sorting and single unit classification*) were included in LMT analyses. For each behavioral session, spike counts were taken in 500-ms bins and only during running periods (> 5 cm/s). Bins with spikes from fewer than 5 cells were discarded. The LMT model assumes that neurons fire according to a Poisson process, and the observed Poisson spikes are governed by temporally smoothed latent variables and a set of nonlinear tuning curves, which are smoothly tuned to the latent variables. This model can simultaneously extract the latent dynamics and tuning curves in an unsupervised manner. The nonlinear tuning curves thus provide the mapping between the low-dimensional latent structure (i.e., neural manifold) and high-dimensional spiking responses. The dimensionality of the latent structure (i.e., number of latent variables) was estimated by calculating the predictive log likelihood via 10-fold cross-validation.

To detect theta sweeps in the latent maps (Fig. 5I), we applied the same Bayesian decoding algorithm as in *Decoding theta and replay sweeps with Bayesian reconstruction*, but we replaced the spatial tuning curve $f(\boldsymbol{x})$ with the latent tuning curve $g(\boldsymbol{l})$, $\boldsymbol{l}$ is a set of latent variables estimated from the LMT model.

**EI ratio**

The excitation-inhibition (EI) ratio (Fig. 3E) was measured as the firing rate ratio between CA1 pyramidal cells (PYRs) and interneurons (INTs). Specifically, we calculated the population spiking activity of all putative PYRs and fast-spiking INTs in CA1. Spike times were binned at 1 ms resolution, and the resulting spike count traces were smoothed using a Gaussian kernel (SD = 15 ms). The EI ratio was then defined as the ratio of smoothed firing rates, as $rate_{PYR}/rate_{INT}$. To assess theta modulation of the EI ratio, we computed the cumulative EI ratio within each azimuthal bin (51 bins), normalized by the total time spent in that bin during either goal-directed sweeps or lateral sweeps. The resulting theta-modulated EI ratio profile was circularly smoothed using a Gaussian kernel (SD = 2.5 bins).

**Putative monosynaptic connection analyses**

Cross-correlograms (CCGs) between pairs of PYR and INT neurons were constructed (Fig. 3, F and G), and only connections from PYR to INT were considered. For visualization, CCGs were rate normalized (i.e., counts/number of reference spikes/$\Delta t$) in units of spikes per sec (Fig. 3F). Further, we identified putative monosynaptic connections of PYR-INT cell pairs by short-lag (2-3 ms) peaks in their CCGs as previously described (*34*, *35*). The peak in the CCG needed to exceed that from the slowly co-modulated baseline, and the peak in the causal direction (positive lags) needed to be significantly larger than the largest peak in the anti-causal direction (negative lags) for further analysis (*34*). To assess the effective strength of PYR-INT synaptic coupling, we calculated spike transmission probability of the PYR-INT cell pairs that exhibited monosynaptic connectivity. To calculate spike transmission probability, the lower frequency baseline $\lambda_{slow}$ was estimated as previously described (*34*, *35*), and the spike transmission probability after N presynaptic spikes using raw CCG normalized by the number of reference spikes was defined as:

$$spike\ transmission\ probability = \frac{\sum_{\tau=0.8ms}^{2.8ms}\left(CCG(\tau)_{observed} - \lambda_{slow}(\tau)\right)}{N}$$

The spike transmission probabilities were further calculated and compared using the CCGs constructed from spikes occurring during goal-directed and lateral theta sweeps (Fig. 3G).

**Simulation of 2D theta sweeps**

We adapted a single-bump continuous attractor network (CAN) model, developed in a previous study (*15*) for modeling left-right alternating theta sweeps (Fig. 3, A-D). In brief (for a more detailed description, see (*15*)), the instantaneous firing rate for a given cell in the network was $f(\boldsymbol{I}(t))$, where $f$ is a rectified function, and $\boldsymbol{I}(t)$ was the input to the cell. Cells fired spikes according to a Poisson process based on $f(\boldsymbol{I}(t))$. The inputs to the cells were a combination of internal ($\boldsymbol{I}^{int}(t)$) and externally driven place input ($\boldsymbol{I}^{P}(t)$), each modulated at theta frequency.

The internal input was given by four components as $\boldsymbol{I}^{int}(t) = \boldsymbol{I}^{rec}(t) + \boldsymbol{I}^{adapt}(t) + \boldsymbol{I}^{goal}(t) + \boldsymbol{I}^{0}$, where $\boldsymbol{I}^{rec}(t)$ is the recurrent input derived from other cells, $\boldsymbol{I}^{adapt}(t)$ is the adaptive inhibitory input, $\boldsymbol{I}^{goal}(t)$ is the goal-direction signal, and $\boldsymbol{I}^{0}$ is the small positive constant bias common to all cells. The place input $\boldsymbol{I}^{P}(t)$ into the cell was given by a 2D Gaussian centered on the rat's position $x_{rat}(t)$. Both internal and place inputs were modulated at theta frequency. The $\boldsymbol{I}^{int}(t)$ was active during the late half of theta cycles (i.e., excitation strength during the late theta phases), and the $\boldsymbol{I}^{P}(t)$ was active during the center 20% of theta cycles, which allows the place inputs to occur briefly around the time of the release of the internally driven dynamics.

**Statistical Analyses**

Data analysis was performed using custom routines in Python, MATLAB (MathWorks) and GraphPad Prism 10 (GraphPad Software). No specific analysis was used to estimate minimal population sample or group size, but the number of animals, sessions and recorded cells was larger or similar to those employed in previous related work (*10*, *14*, *19*, *22–24*). Unless otherwise noted, the non-parametric Wilcoxon rank-sum or Wilcoxon signed-rank test was used for unpaired and paired data comparisons, respectively, and ANOVA and Friedman test were used for multiple comparisons. All statistical tests were two-tailed with $p < 0.05$ as the cutoff for statistical significance, which is indicated by asterisks (*$p$<0.05, **$p$<0.01, ***$p$<0.001, and ****$p$<0.0001). Error bars show SEMs, and boxplots show median (central mark), 75th (box), and 90th (whiskers) percentile, unless indicated otherwise.

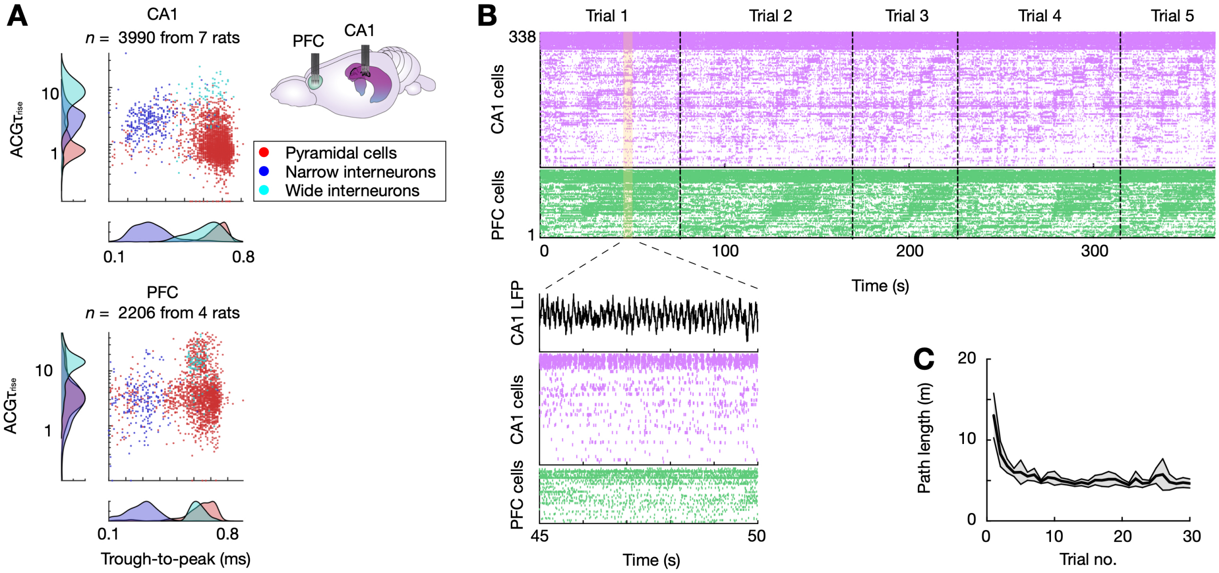


**Figure S1. Simultaneous CA1 – PFC silicon probe recordings in behaving rats.**

**(A)** Cell-type classification for CA1 (*top*) and PFC neurons (*bottom*), showing spike waveform width (trough-to-peak) and the temporal scale of the rising phase of the auto-correlograms (ACGs). Diagram on the right illustrate probe implant. **(B)** Example of representative recording. Raster plot of CA1 (purple) and PFC (green) neurons during five consecutive trials in the goal-directed navigation (*gdn*) task, sorted by seqNMF (*68*) (corresponding interneurons shown on top). Inset below shows a 5-second fragment with CA1 LFP trace on top (note theta oscillations). **(C)** Learning performance in the *gdn* task quantified as path length to retrieve the three hidden water rewards ($n$ = 26 novel sessions in 5 rats, mean ± 95% CI).

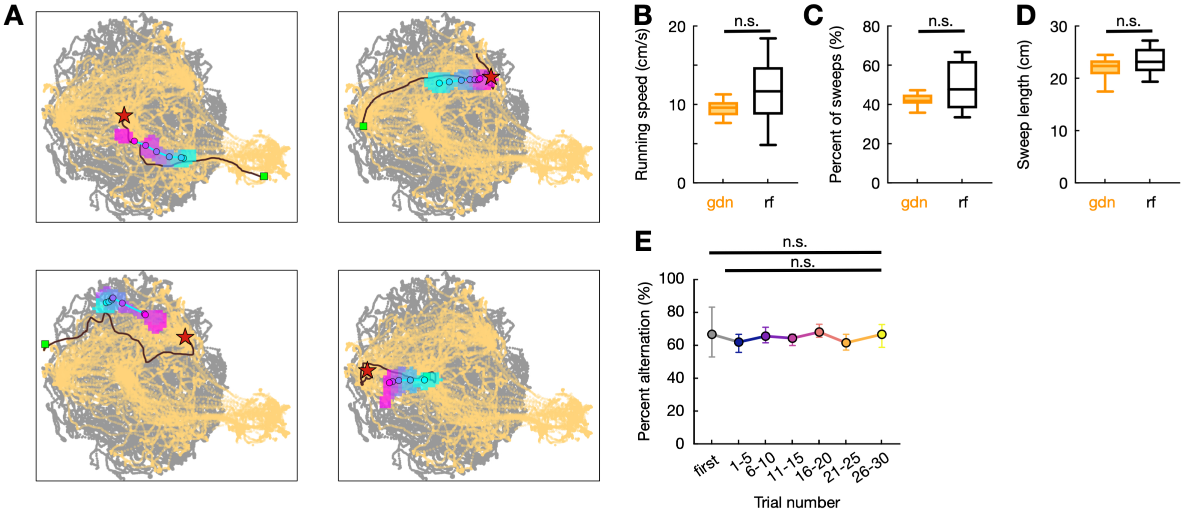


**Figure S2. Additional examples and quantification of theta sweeps.**

**(A)** Four additional examples of goal-directed theta sweeps. Gray and yellow traces: positions visited during *rf* and *gdn tasks*, respectively; black lines: current trajectory; green square: trajectory start; red star: next chosen goal; colored blobs: decoded positions, colored by time within sweep. **(B)** Average running speed during *rf* and *gnd* tasks (*rf*: $n = 12$ sessions, 6 rats; *gdn*: $n = 63$ sessions, 5 rats; $p = 0.076$, rank-sum test). **(C)** Average sweep length during *rf* and *gdn tasks* ($p = 0.32$, rank-sum test). **(D)** Percentage of significant theta sweeps in theta cycles during *rf* and *gdn tasks* ($p = 0.20$, rank-sum test). **(E)** Percentage of alternating sweeps across trials in the *gdn* tasks (mean ± 95% CI; $p = 0.74$, one way ANOVA).

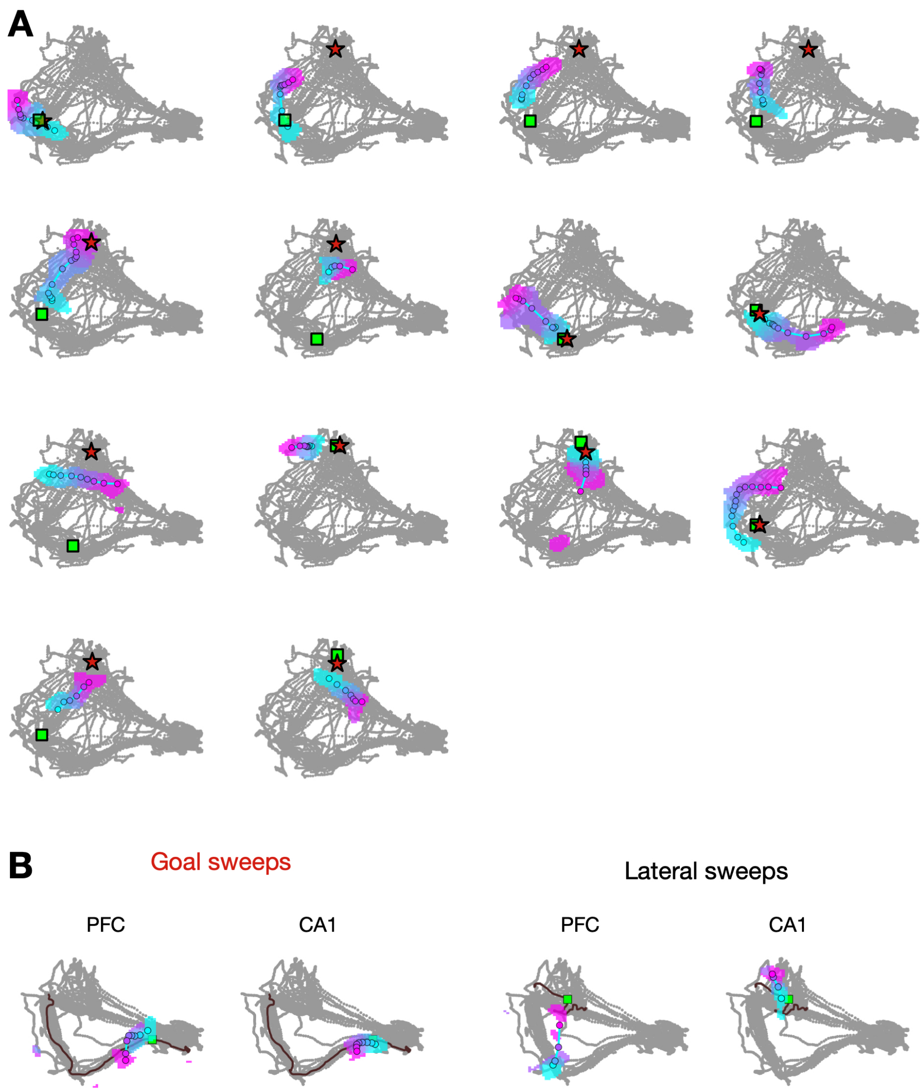


**Figure S3. Additional examples of replay sweeps and CA1-PFC coordination during theta sweeps.**

**(A)** All replay events that occurred on the maze during the representative session of the *gdn* task shown in Fig. 4A. red stars: closer goal location to replay decoded trajectory; green square: current animal position; colored blob: decoded position over replay time; filled circles: positions with maximal decoded probability; gray lines: animal trajectories in the whole session). **(B)** Additional examples of simultaneously decoded positions in PFC and CA1 during a hippocampal goal-directed theta sweep (*left*) and a lateral sweep (*right*). Data is plotted as in Fig. 4, E and F. Note the similarity between PFC and CA1 decoded spatial trajectory during goal-directed but not lateral theta sweep.

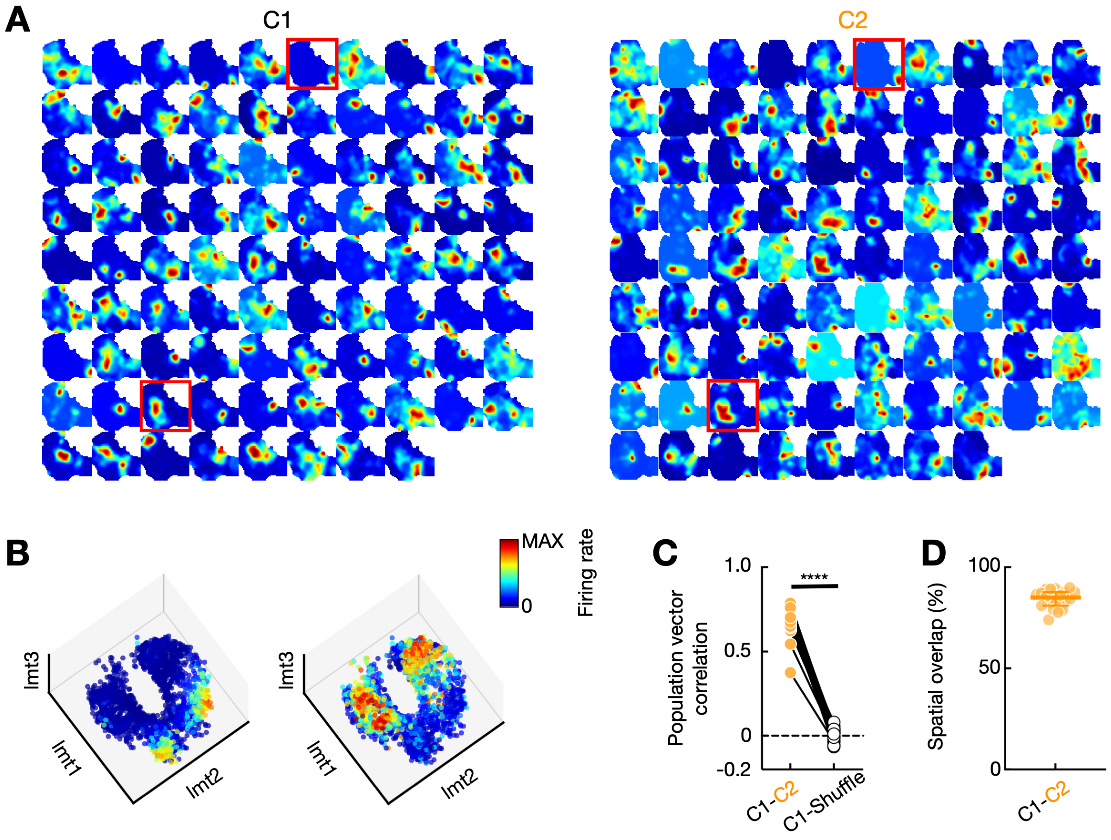


**Figure S4. Additional control analyses for quantifying the latent maps in the hippocampus.**

**(A)** Spatial rate maps of simultaneously recorded CA1 place cells in the *gdn* task with two different reward configurations (C1 and C2). Red squares denote the two example cells shown in **B**. **(B)** Tuning curves of three latent variables ($lmt_1$–$lmt_3$) inferred by the LMT model for the two example cells shown in **A**. Each dot represents a latent state at time $t$, given by $[lmt_1(t), lmt_2(t), lmt_3(t)]$, with color denoting the cell's firing rate at that state. **(C)** Population vector correlation between the two configurations, compared to cell-identity shuffles (****$p$ = 3.81e-16, Wilcoxon paired test, $n$ = 19 C1–C2 session pairs from 5 rats). **(D)** Spatial overlap, quantified as the percentage of spatial bins visited by the animal in both configurations ($n$ = 19 C1–C2 session pairs from 5 rats).

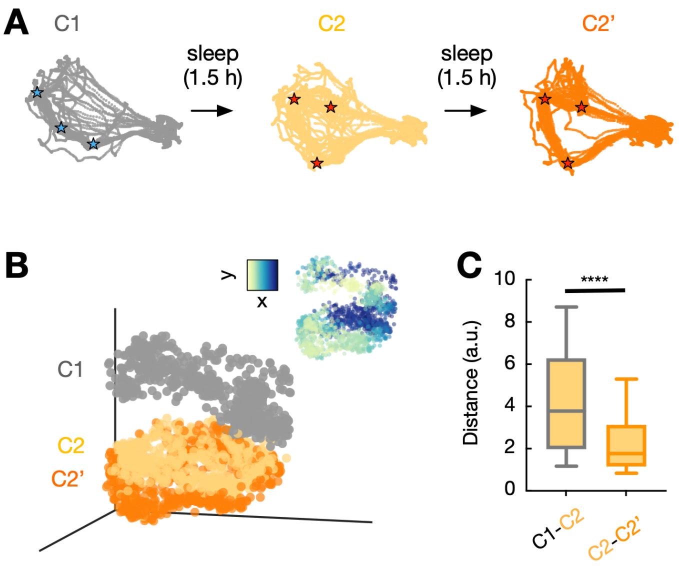


**Figure S5. Latent map separation cannot be attributed solely to temporal drift.**

**(A)** Task design. In a subset of experiments ($n$ = 6 days from 4 rats), animals performed three 20-trial sessions in either the C1–C1'–C2 or C1–C2–C2' order, with ~1.5 hours of sleep interleaved between sessions. **(B)** Representative sessions demonstrate that the latent map for one configuration is largely separated from that of a different configuration, while the separation between a configuration and its re-exposure is modest. **(C)** Latent map separation quantified as the Euclidean distance between latent states from two sessions (****$p < 1e{-}16$, Wilcoxon paired test, $n$ = 6 days from 4 rats).

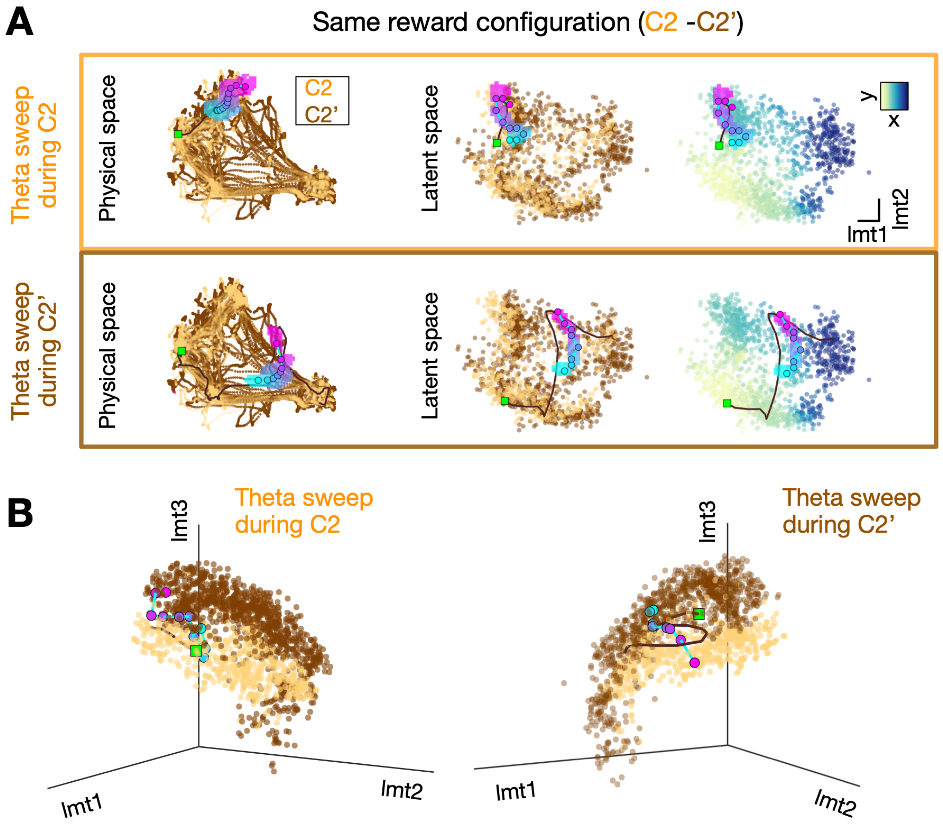


**Figure S6. Two example theta sweeps on C2 and C2' latent maps.**

**(A)** *Left*, theta sweeps shown in physical space, as in Fig. 5I (orange for C2; brown for re-exposure to C2, or C2'). *Middle* and *Right*, the same theta sweeps visualized in the 2D latent space estimated by the LMT model, color-coded by configuration (*middle*) and by location (*right*). **(B)** The two example theta sweeps plotted in the full 3D latent space.

**Movie S1. Emergence of goal-directed theta sweeps via reduced feedback inhibition and an egocentric goal signal.**

This video shows a model simulation of theta sweeps during both random foraging and goal-directed navigation (see also Fig. 3B). Each theta sweep is represented by a time-colored line, with the black line indicating the simulated rat's trajectory and the red star marking the goal location. During random foraging, left-right alternating sweeps are observed. In contrast, goal-directed sweeps emerge when an egocentric goal signal is present (*top left*) and feedback inhibition is reduced during the late phase of theta (*top right*).
